# ACE2-expressing endothelial cells in aging mouse brain

**DOI:** 10.1101/2020.07.11.198770

**Authors:** Su Bin Lim, Valina L. Dawson, Ted M. Dawson, Sung-Ung Kang

## Abstract

Angiotensin-converting enzyme 2 (ACE2) is a key receptor mediating the entry of SARS-CoV-2 into the host cell. Through a systematic analysis of publicly available mouse brain sc/snRNA-seq data, we found that ACE2 is specifically expressed in small sub-populations of endothelial cells and mural cells, namely pericytes and vascular smooth muscle cells. Further, functional changes in viral mRNA transcription and replication, and impaired blood-brain barrier regulation were most prominently implicated in the aged, ACE2-expressing endothelial cells, when compared to the young adult mouse brains. Concordant EC transcriptomic changes were further found in normal aged human brains. Overall, this work reveals an outline of ACE2 distribution in the mouse brain and identify putative brain host cells that may underlie the selective susceptibility of the aging brain to viral infection.

## Introduction

In addition to the well-known respiratory symptoms, COVID-19 patients suffer from a loss of smell and taste, headache, impaired consciousness, and nerve pain [1], raising possibility of virus infiltration in the nervous system, including brain. Despite cases emerging of COVID-19 patients with neurologic manifestations, potential neurotropic mechanisms underlying SARS-CoV-2-mediated entry into the cells of the brain are largely unexplored.

Evidenced by transgenic mice models [2,3], the evolutionarily-related coronaviruses, such as SARS-CoV and MERS-CoV, can invade the brain by replicating and spreading through the nasal cavity, and possibly olfactory bulbs located in close proximity to the frontal lobes of the brain [4]. Once inside the brain, viruses can harm the brain directly and indirectly by infecting the cells and myelin sheaths, and by activating microglia, which may in turn consume healthy neurons to induce neuroinflammation and neurodegeneration [5].

Many of the observed neurological symptoms observed may in part be explained by a primary vasculopathy and hypercoagulability [6], as endothelial dysfunction and the resulting clotting are increasingly being observed in patients with severe COVID-19 infection [7–9]. Consistent with these findings, the first pathologic evidence of direct viral infection of the EC and lymphocytic endotheliitis has been found in multiple organs, including lung, heart, kidney and liver, in a series of COVID-19 patients [10].

In contrast, recent clinical findings, including an MRI study [11] and immunohistochemistry and RT-qPCR analyses [12,13], did not observe any signs of encephalitis from postmortem brain examination of COVID-19 patients. Similarly, postmortem analysis of SARS-CoV-2-exposed mice transgenically expressing ACE2 via mouse ACE2 promoter failed to detect the virus in the brain [14]. In light of such controversy regarding neuropathological features, a more comprehensive assessment on the distribution of ACE2 in a cell type-specific manner is required to identify putative brain host cells.

Here, we analyzed publicly available and spatially rich brain RNA-seq datasets to assess ACE2 distribution in mouse brains at the single-cell and single-nuclei level. We found that ACE2 was consistently expressed in small subpopulations of endothelial cells (ECs) and mural cells in all the analyzed datasets, in which the impaired blood-brain barrier was further implicated in the aged brains. These findings altogether may hold potential to initiate new avenues of research on specific cells types (EC and vascular mural cells) that remain poorly understood particularly in relation to the aging and viral infection in the brain.

## Results

### ACE2 is expressed in small sub-populations of ECs and mural cells in the adult mouse brain

A total of 11 adult mouse brains datasets deposited in the Single Cell Portal (SCP) were analyzed in this study (**Table 1**; see *Methods*). Datasets lacking vascular cell types were excluded from the data collection and subsequent analyses. The analyzed datasets were derived from diverse single-cell (scRNA-seq) and single-nuclei (snRNA-seq) sequencing technologies, including 10x Genomics, SMART-seq, Drop-seq, and sNucDrop-seq. The number of cells included in the final SCP-generated tSNE/UMAP plots also varied from 1,301 (lowest; T3) to 611,034 (largest; T11).

**Table 1.** Mouse brain sc/snRNA-seq datasets analyzed in this study.

| Dataset notation | No. of cells | Region | Sequencing technology | Study name in SCP |
| --- | --- | --- | --- | --- |
| Single-cell RNA sequencing (scRNA-seq) |  |  |  |  |
| T1 | 7,871 | Cortex (auditory cortex) | * | Mouse Auditory Cortex Sample |
| T2 | 1,301 | Cortex (anterior lateral motor cortex) | SMART-seq | A transcriptomic taxonomy of adult mouse anterior lateral motor cortex (ALM) |
| T3 | 72,543 | Vascular cells | 10x Genomics | Aging_mouse_brain_kolab |
| T4 | 20,921 | Hypothalamus | Drop-seq | A molecular census of arcuate hypothalamus and median eminence cell types |
| T5 | 1,679 | Cortex (primary visual cortex) | SMART-seq | A transcriptomic taxonomy of adult mouse visual cortex (VISp) |
| T6 | 37,069 | Whole brain | 10x Genomics | Aging Mouse Brain |
| Single-nucleus RNA sequencing (snRNA-seq) |  |  |  |  |
| T7 | 3,402 | Cortex | 10x Genomics | Experiment 2 Mouse PBS |
| T8 | 13,861 | Midbrain (SNr, SNc, VTA) | 10x Genomics | Single nuclei dataset, SN/VTA (MD720) |
| T9 | 13,783 | Cortex | Multiple<sup>†</sup> | Single Cell Comparison: Cortex data |
| T10 | 611,034 | Cerebellum | 10x Genomics | A transcriptomic atlas of the mouse cerebellum |
| T11 | 18,194 | Cortex | sNucDrop-seq | sNucDrop-seq: Dissecting Cell-Type Composition and Activity-Dependent Transcriptional State in Mammalian Brains by Massively Parallel Single-Nucleus RNA-Seq |

\* Integrated data of the Brain Architecture Portal and the Single Cell Portal
(https://singlecell.broadinstitute.org/single_cell/study/SCP280/mouse-auditory-cortex-sample#study-summary)
<sup>†</sup> Smart-seq2, 10x Chromium, Drop-seq, and sci-RNA-seq

Despite the varying library preparation and sequencing technologies, which arguably recover population heterogeneity to different extents, the analysis of the retrieved scRNA-seq datasets (**Fig. 1 *A*-*E***) consistently showed increased levels of ACE2 in a small subpopulation of vascular cells, namely endothelial cells (EC), pericytes (PC), and vascular smooth muscle cells (VSMC), across different brain regions, including auditory cortex (T1), anterior lateral motor cortex (T2), primary visual cortex (T5), hypothalamus (T4), and whole brain (T6). Further, scRNA-seq dataset specifically derived from brain vasculature in young adult and aged mice (T3) confirms the elevated ACE2 expression in subsets of the three identified cell types, which consist of 32.8% of the cell populations (**Fig. 1 *C***). Similarly, ACE2 mRNAs were enriched in subpopulations of ECs and mural cells of all analyzed snRNA-seq datasets (**Fig. 2 *A*-*E***) derived from cortex (T7, T9, T11), midbrain (T8), including substantia nigra pars reticulata (SNr), substantia nigra pars compacta (SNc), and ventral tegmental area (VTA), and cerebellum (T10).

**Fig. 1.**
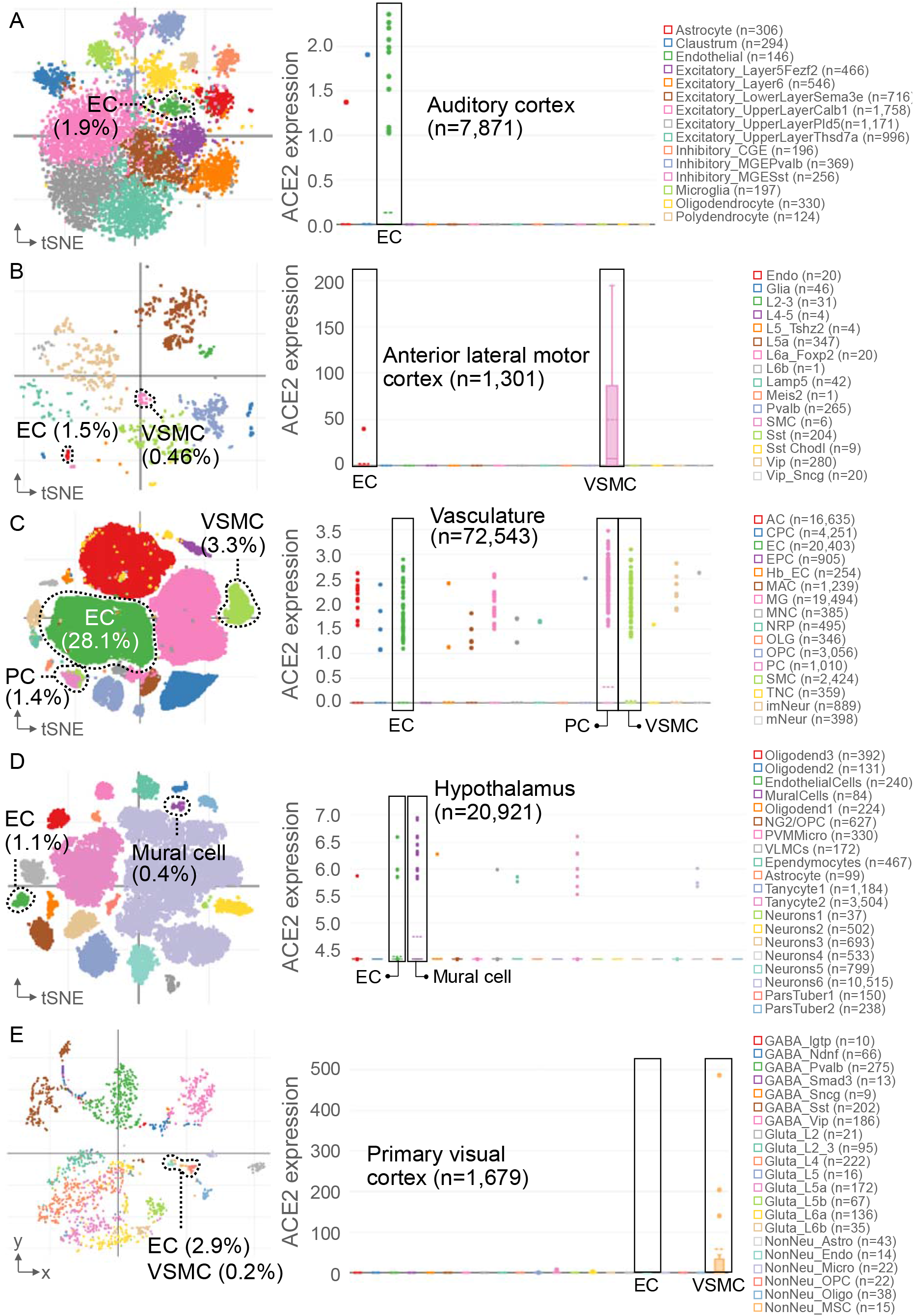
ACE2 expression in SCP scRNA-seq datasets. Brain region-specific t-SNE or UMAP visualization (left) and boxplots (including outliers) of cell type-specific ACE2 expression (right) for Dataset (A) T1, (B) T2, (C) T3, (D) T4, and (E) T5. The percentage of EC, PC and VSMC cell populations are shown. n indicates the total number of cells included in the cell clustering.

**Fig. 2.**
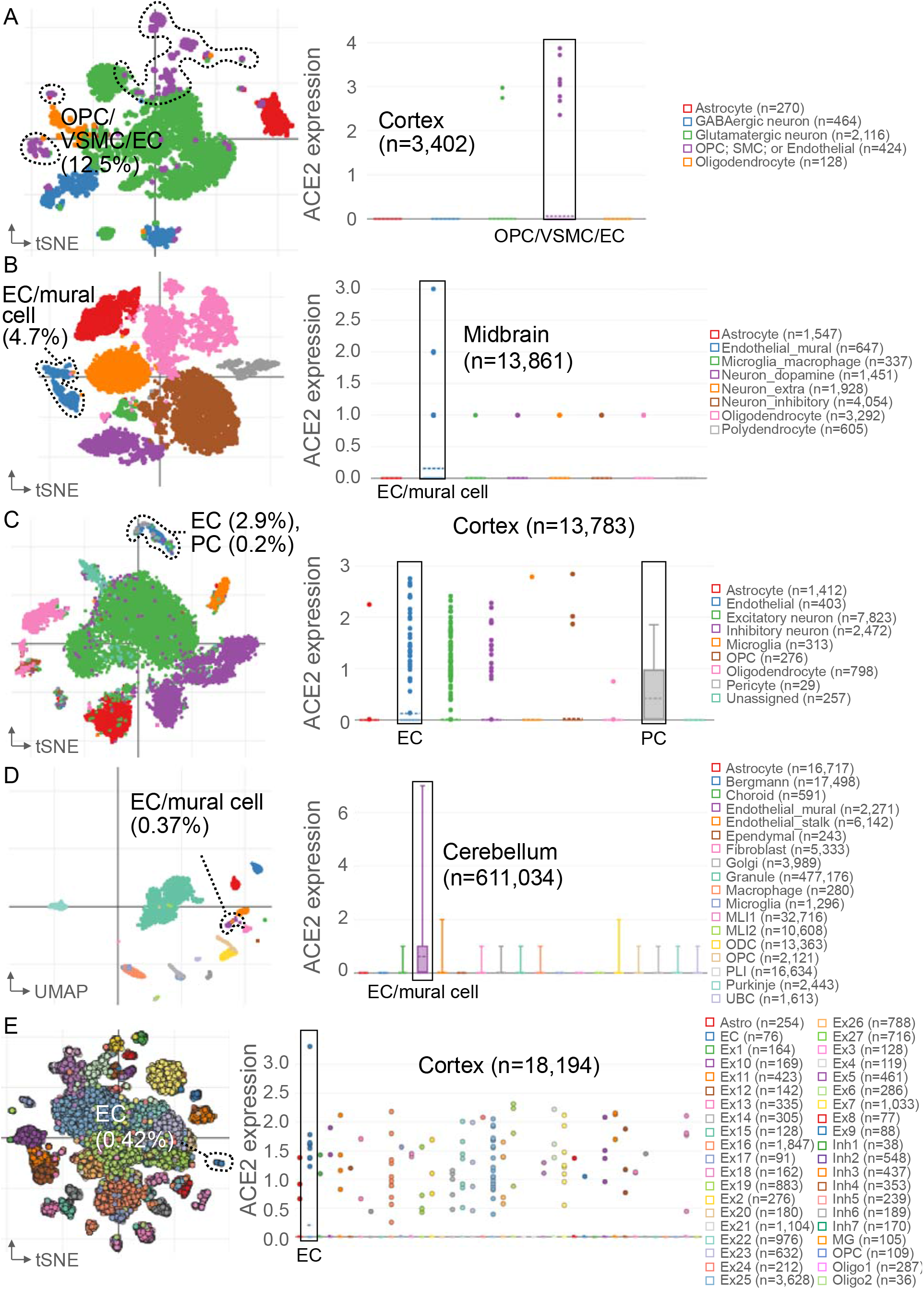
ACE2 expression in SCP snRNA-seq datasets. Brain region-specific t-SNE or UMAP visualization (left) and boxplots (including outliers) of cell type-specific ACE2 expression (right) for Dataset (A) T7, (B) T8, (C) T9, (D) T10, and (E) T11. The percentage of EC, PC and VSMC cell populations are shown. n indicates the total number of cells included in the cell clustering.

### Impaired blood-brain barrier is implicated in ECs of the aged brain

We next asked if these identified vascular cell sub-populations expressing ACE2 would be affected by aging and whether they have unique transcriptional changes that are functionally important. Of the 25 major cell types of different lineages (oligodendrocyte, astrocyte, and neuronal lineages, ependymal cells, vasculature cells, and immune cells), EC, PC, and VSMC cell types contribute to a majority of ACE2-expressing cells in the whole mouse brain (T6) (**Fig. 3 *A*-*C***). tSNE plots (**Fig. 3 *A*-*B***) show projection of 37,069 single cells (16,028 young and 21,041 old) derived from the brains of 8 young (2-3 months) and 8 old (21-23 months) mice.

**Fig. 3.**
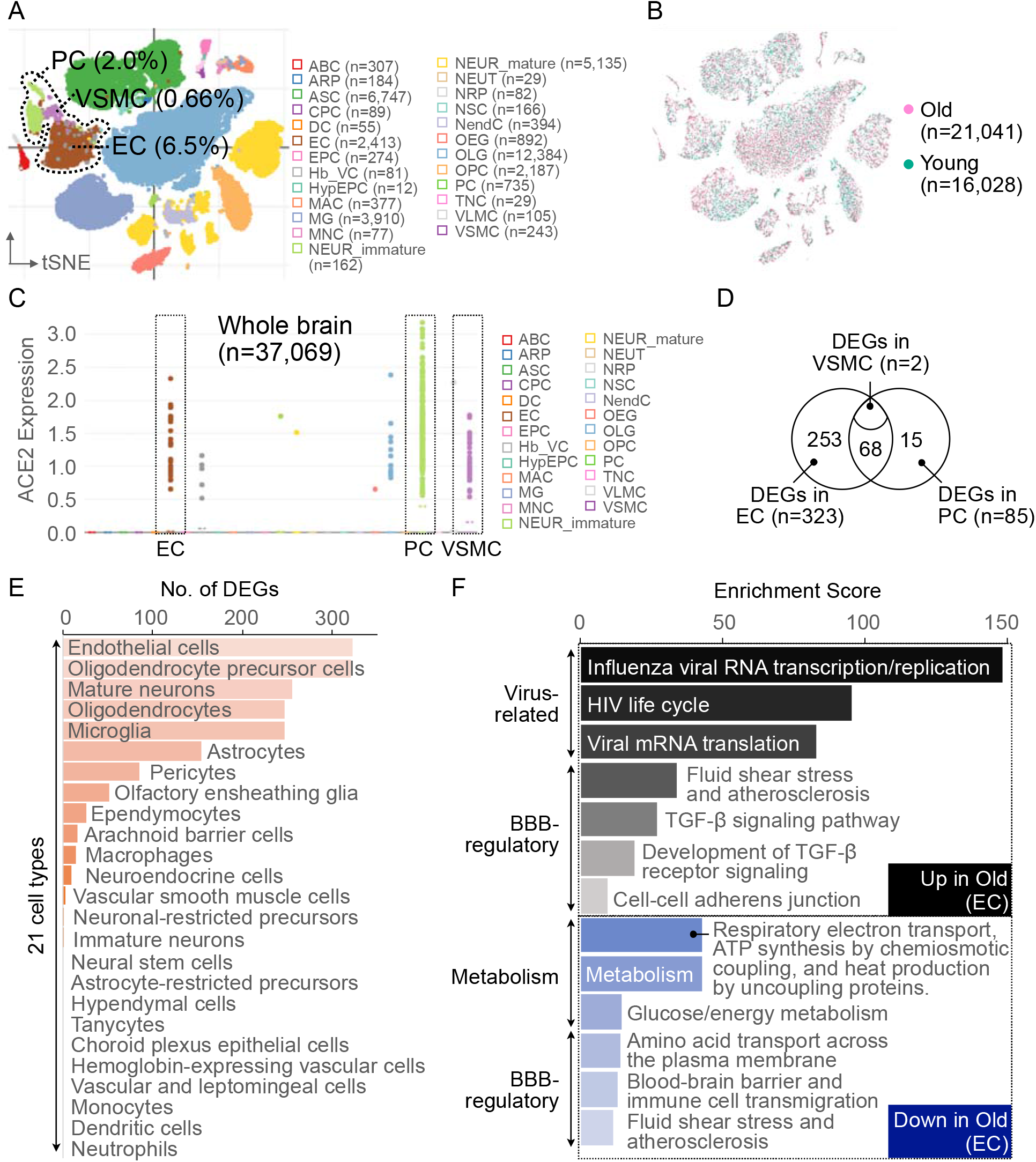
Impaired BBB implicated in ACE2-expressing ECs of the aged brain. (A) t-SNE visualization of the aging mouse brain colored by cell type (*left*) and age (*right*). The percentage of EC, PC and VSMC cell populations are shown. (B) Cell type-specific ACE2 expression. n indicates the total number of cells included in the cell clustering. (C) Cell types ranked by the number of DEGs (young vs. old) and Venn Diagram depicting the overlapping DEGs in EC, PC and VSMC cell populations. (D) Pathway enrichment analysis (GeneAnalytics) of the EC-specific DEGs.

Importantly, we found that the EC was the most differentially expressed (DE) cell type in the aged mouse brain, as compared to the young, in which a set of 68 DE genes (DEGs) was further concordantly up- and down-regulated in the aged PCs (**Fig. 3 *D*-*E*;** see *Methods*). Given that PCs are in direct contact with ECs, covering between 22% and 99% of the EC surface [15], such shared transcriptomic changes highlight defective pericyte-endothelial interface as the most notable change occurring due to aging in mouse brains. Notably, viral RNA transcription/translation, blood-brain barrier (BBB) regulation and glucose/energy metabolism were among the most prominently enriched functional pathways in the ECs of the aged mouse brain (**Fig. 3 *F***; see *Methods*). These findings altogether indicate that the pericyte-endothelial interface may serve as a potential target and reservoir of virus in the brain, with increased susceptibility to its infection in the aged brain with BBB defects.

### Aged mouse brain EC transcriptomic signatures are enriched in aged human brains

To assess our findings in a human context, we next asked if the identified transcriptomic changes in the aged EC gene signatures would further be conserved and detected in normal human aged brains using bulk RNA-seq data derived from the Genotype-Tissue Expression (GTEx) project database (**Table 2**) [16]. Expression levels (in TPM) of human orthologs of the aged EC DEGs matched in the GTEx data were compared between the young (< 60) and old (≥ 60 yrs) human samples (see *Methods*). Of all genes evaluated (n = 56,200), 8,215 (15%) showed the expression level significantly different (t-test q-value ≤ 0.05; see *Methods*) while the expression levels of 37% of EC DEGs were different in the old group from that of the young population, indicating concordant EC transcriptomic changes in normal aged human brains (**Fig. 4 *A*-*B***). This conclusion was further supported by Gene Set Enrichment Analysis (GSEA) results, displaying a significant enrichment of the aged (n = 141) and young (n = 133) mouse EC DEGs in the old (n = 814) and young (n = 621) human samples, respectively (**Fig. 4 *C***). In conclusion, we have identified specific EC signatures that are functionally important and related to the aging and viral infection in the brain.

**Fig. 4.**
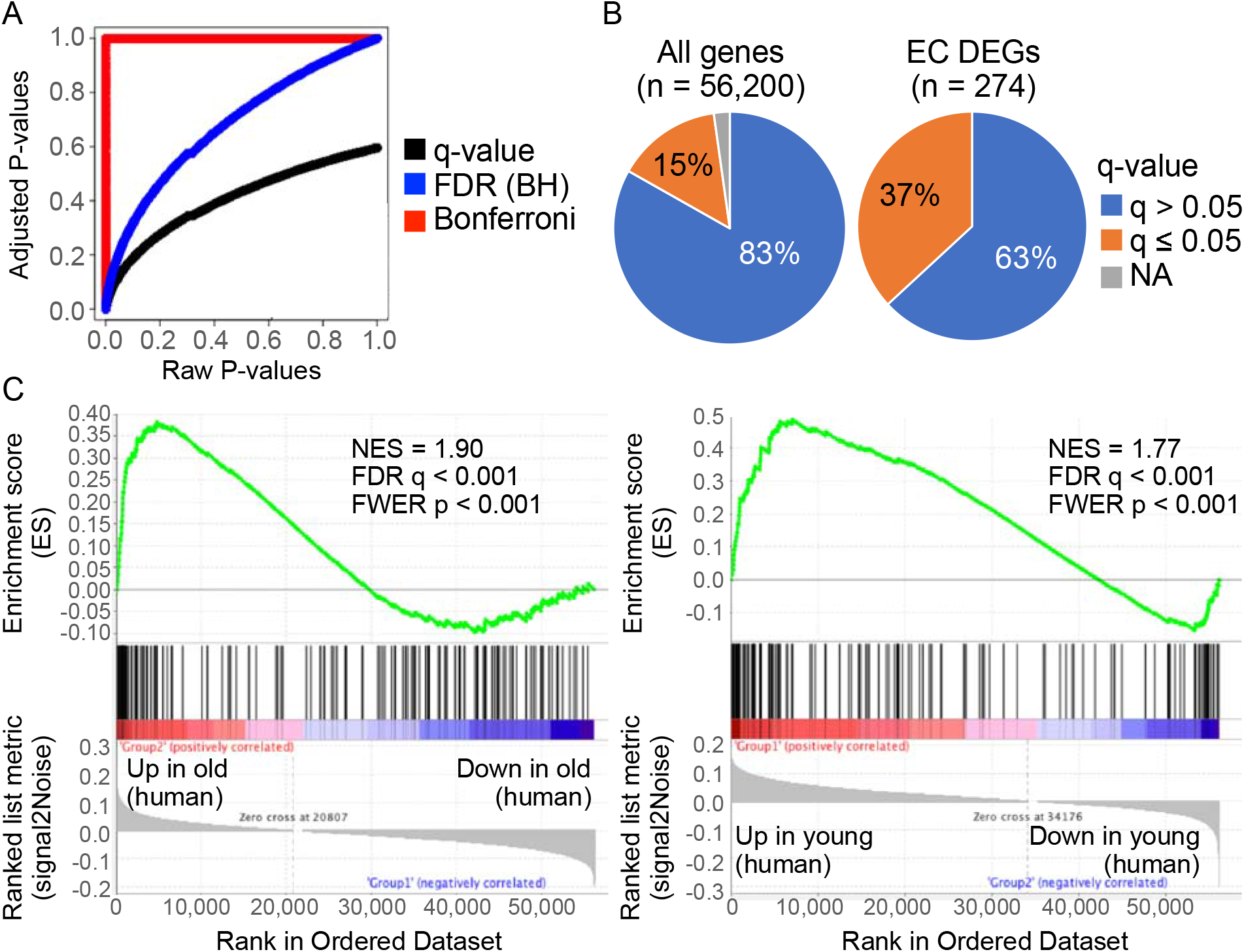
Aged mouse brain EC DEGs assessed in human brain bulk RNA-seq samples (GTEx). (A) Adjusted P-values via three different correction methods and their relationship to raw P-values (*t*-test). (B) Pie charts depicting the proportion (%) of all genes (*left*) and EC DEGs (*right*) (C) GSEA-generated enrichment plots for the aged (left) and the young mouse EC DEGs (right) gene sets. NES = normalized enrichment score; FDR = false discovery rate; FWER = family-wise error rate.

**Table 2.** Human brain bulk RNA-seq data (GTEx) analyzed in this study.

| Brain region | # of total samples | # of young (< 60 yrs) | # of old (≥ 60 yrs) | # of female | # of male |
| --- | --- | --- | --- | --- | --- |
| Amygdala | 88 | 39 | 49 | 25 | 63 |
| Anterior cingulate cortex | 107 | 47 | 60 | 30 | 77 |
| Caudate (basal ganglia) | 142 | 65 | 77 | 35 | 107 |
| Cerebellar hemisphere | 121 | 54 | 67 | 34 | 87 |
| Cerebellum | 138 | 62 | 76 | 35 | 103 |
| Hippocampus | 123 | 48 | 75 | 34 | 89 |
| Frontal cortex (BA9) | 124 | 50 | 74 | 32 | 92 |
| Cortex | 137 | 65 | 72 | 41 | 96 |
| Hypothalamus | 120 | 47 | 73 | 30 | 90 |
| Nucleus accumbens (basal ganglia) | 140 | 62 | 78 | 34 | 106 |
| Putamen (basal ganglia) | 121 | 55 | 66 | 25 | 96 |
| Substantia nigra | 74 | 27 | 47 | 21 | 53 |
| # of total samples | 1435 | 621 | 814 | 376 | 1059 |

## Discussion

While our study provides a foundation for a more refined level of analysis of EC and vascular PC, a cell type that remains poorly understood despite its key roles in immune response and microvascular stability [17], our analyses are limited only to the normal aging mouse and human brains, lacking the context of COVID-19 neuropathology. A number of recent sc/snRNA-seq studies identified ACE2 mRNA in the olfactory neuroepithelium [18,19], although there are no sc/snRNA-seq data derived from postmortem brains of COVID-19 patients to date. The distribution of ACE2 and other genes mediating SARS-CoV-2 entry into the cells of the brain thus remains to be investigated across different regions and cell types.

Using scRNA-seq data derived from normal human brain tissues, Muus, C *et al.* [20] have identified ACE2^+^TMPRSS2^+^ oligodendrocytes, while Chen *et al.* [21] have found subsets of both neuronal (excitatory and inhibitory neurons) and non-neuronal cells (mainly astrocytes and oligodendrocytes) expressing ACE2. These studies, however, have only used a limited number of datasets, which may in part explain the inconsistent results of the identified cell types.

Despite the works that failed to identify direct signs of SARS-CoV-2 infection in the brains of COVID-19 patients [12,13], other lines of evidence support the neurotropism of the virus, as evidenced by experimental platforms leveraging human induced pluripotent stem cell (iPSC)-derived dopaminergic neurons [22] and an organotypic brain model [23]. At this point, more data and systematic molecular evidence will be needed to assess the neuroinvasive potential of SARS-CoV-2 and its potential impact on neuroinflammation and neurodegenerative diseases.

## Methods

### Mouse brain sc/snRNA-seq data and analysis

Of all the independent studies retrieved with the term “ACE2” from the Single Cell Portal (https://singlecell.broadinstitute.org/single_cell), 11 sc/snRNA-seq datasets were derived from (1) adult (young or old) mouse brains, and had (2) author-defined cell type annotations including endothelial cells (EC, PC, or VSMC). A total of 801,658 cells were analyzed in this study, based on the organ (i.e., brain) and species of origin (i.e., mouse), and diseased status (i.e., normal). 2-D tSNE/UMAP plots (colored by cell type) and box plots for cell type-specific ACE2 expression presented in this study were generated by the Single Cell Portal. t-SNE visualization (colored by age) and 25 cell type-specific DE gene lists of the aging mouse brains (T1) were obtained from the advanced interactive data viewer (http://shiny.baderlab.org/AgingMouseBrain/AgingMouseBrain_SCV/). Genes with |logGER (Gene Expression Ratio)|>0.1 and FDR<0.05 were defined as DEGs, and were analyzed for functional pathway enrichment using GeneAnalyics (https://geneanalytics.genecards.org/).

### Human brain bulk RNA-seq data and analysis

De-identified processed human brain bulk RNA-seq data and annotation files for sample attributes and subject phenotypes were obtained from the Genotype-Tissue Expression (GTEx) portal (https://www.gtexportal.org/home/datasets). Data from amygdala (*n* = 88), anterior cingulate cortex (*n* = 107), caudate (basal ganglia) (*n* = 142), cerebellar hemisphere (*n* = 121), cerebellum (*n* = 138), hippocampus (*n* = 123), frontal cortex (BA9) (*n* = 124), cortex (*n* = 137), hypothalamus (*n* = 120), nucleus accumbens (basal ganglia) (*n* = 140), putamen (basal ganglia) (*n* = 121), substantia nigra (*n* = 74) were analyzed in this study. A total of 1,435 samples were divided into two age groups: young (< 60) and old (≥ 60 years old). Expression levels (in TPM) were compared between the two age groups for all genes (*n* = 56,200) by *t*-test with multiple testing correction. The performance of Bonferroni correction and False Discovery Rate (FDR)-Benjamini-Hochberg (BH) procedure was assessed using the RStudio (Version 1.2.5019) base function p.adjust(). The R *qvalue* package [24] from Bioconductor was used to assess the performance of q-value approach. The R *biomaRt* package [25] from Bioconductor was used to convert mouse EC DEGs to human gene symbols (hgnc) using getLDS() function. Gene Set Enrichment Analysis (GSEA v4.0.3; https://www.gsea-msigdb.org/gsea/index.jsp) [26] was used to assess EC DEGs (EC up- and down-regulated genes) in GTEx human brain bulk RNA-seq samples. Following parameters were set to run enrichment tests: (1) number of permutations = 1,000, (2) collapse/remap to gene symbols = no_collapse, and (3) permutation type = gene_set.

## Acknowledgements

T.M.D. is the Leonard and Madlyn Abramson Professor in Neurodegenerative Diseases.

## Author contributions

V.L.D., T.M.D., and S.U.K. supervised the project; S.B.L., V.L.D., T.M.D. and S.U.K. formulated the hypothesis; S.B.L. and S.U.K. performed research; S.B.L., V.L.D., T.M.D., and S.U.K. wrote the paper.

## Reference

1. Mao, L.; Jin, H.; Wang, M.; Hu, Y.; Chen, S.; He, Q.; Chang, J.; Hong, C.; Zhou, Y.; Wang, D., et al. Neurologic Manifestations of Hospitalized Patients With Coronavirus Disease 2019 in Wuhan, China. JAMA neurology 2020, 10.1001/jamaneurol.2020.1127, doi:10.1001/jamaneurol.2020.1127.

2. Netland, J.; Meyerholz, D.K.; Moore, S.; Cassell, M.; Perlman, S. Severe acute respiratory syndrome coronavirus infection causes neuronal death in the absence of encephalitis in mice transgenic for human ACE2. Journal of virology 2008, 82, 7264–7275, doi:10.1128/JVI.00737-08.

3. Li, K.; Wohlford-Lenane, C.; Perlman, S.; Zhao, J.; Jewell, A.K.; Reznikov, L.R.; Gibson-Corley, K.N.; Meyerholz, D.K.; McCray, P.B., Jr. Middle East Respiratory Syndrome Coronavirus Causes Multiple Organ Damage and Lethal Disease in Mice Transgenic for Human Dipeptidyl Peptidase 4. The Journal of infectious diseases 2016, 213, 712–722, doi:10.1093/infdis/jiv499.

4. Li, Y.-C.; Bai, W.-Z.; Hashikawa, T. The neuroinvasive potential of SARS-CoV2 may play a role in the respiratory failure of COVID-19 patients. J Med Virol 2020, 92, 552–555, doi:10.1002/jmv.25728.

5. Jang, H.; Boltz, D.; Sturm-Ramirez, K.; Shepherd, K.R.; Jiao, Y.; Webster, R.; Smeyne, R.J. Highly pathogenic H5N1 influenza virus can enter the central nervous system and induce neuroinflammation and neurodegeneration. Proceedings of the National Academy of Sciences 2009, 106, 14063, doi:10.1073/pnas.0900096106.

6. Cooper, K.W.; Brann, D.H.; Farruggia, M.C.; Bhutani, S.; Pellegrino, R.; Tsukahara, T.; Weinreb, C.; Joseph, P.V.; Larson, E.D.; Parma, V., et al. COVID-19 and the Chemical Senses: Supporting Players Take Center Stage. Neuron 2020, 10.1016/j.neuron.2020.06.032, doi:10.1016/j.neuron.2020.06.032.

7. Klok, F.A.; Kruip, M.; van der Meer, N.J.M.; Arbous, M.S.; Gommers, D.; Kant, K.M.; Kaptein, F.H.J.; van Paassen, J.; Stals, M.A.M.; Huisman, M.V., et al. Incidence of thrombotic complications in critically ill ICU patients with COVID-19. Thrombosis research 2020, 191, 145–147, doi:10.1016/j.thromres.2020.04.013.

8. Zhou, F.; Yu, T.; Du, R.; Fan, G.; Liu, Y.; Liu, Z.; Xiang, J.; Wang, Y.; Song, B.; Gu, X., et al. Clinical course and risk factors for mortality of adult inpatients with COVID-19 in Wuhan, China: a retrospective cohort study. Lancet 2020, 395, 1054–1062, doi:10.1016/S0140-6736(20)30566-3.

9. Zhang, Y.; Xiao, M.; Zhang, S.; Xia, P.; Cao, W.; Jiang, W.; Chen, H.; Ding, X.; Zhao, H.; Zhang, H., et al. Coagulopathy and Antiphospholipid Antibodies in Patients with Covid-19. The New England journal of medicine 2020, 382, e38, doi:10.1056/NEJMc2007575.

10. Varga, Z.; Flammer, A.J.; Steiger, P.; Haberecker, M.; Andermatt, R.; Zinkernagel, A.S.; Mehra, M.R.; Schuepbach, R.A.; Ruschitzka, F.; Moch, H. Endothelial cell infection and endotheliitis in COVID-19. Lancet 2020, 395, 1417–1418, doi:10.1016/S0140-6736(20)30937-5.

11. Coolen, T.; Lolli, V.; Sadeghi, N.; Rovai, A.; Trotta, N.; Taccone, F.S.; Creteur, J.; Henrard, S.; Goffard, J.C.; De Witte, O., et al. Early postmortem brain MRI findings in COVID-19 non-survivors. Neurology 2020, 10.1212/WNL.0000000000010116, doi:10.1212/WNL.0000000000010116.

12. Schaller, T.; Hirschbuhl, K.; Burkhardt, K.; Braun, G.; Trepel, M.; Markl, B.; Claus, R. Postmortem Examination of Patients With COVID-19. Jama 2020, 10.1001/jama.2020.8907, doi:10.1001/jama.2020.8907.

13. Solomon, I.H.; Normandin, E.; Bhattacharyya, S.; Mukerji, S.S.; Keller, K.; Ali, A.S.; Adams, G.; Hornick, J.L.; Padera, R.F., Jr.; Sabeti, P. Neuropathological Features of Covid-19. The New England journal of medicine 2020, 10.1056/NEJMc2019373, doi:10.1056/NEJMc2019373.

14. Bao, L.; Deng, W.; Huang, B.; Gao, H.; Liu, J.; Ren, L.; Wei, Q.; Yu, P.; Xu, Y.; Qi, F., et al. The pathogenicity of SARS-CoV-2 in hACE2 transgenic mice. Nature 2020, 10.1038/s41586-020-2312-y, doi:10.1038/s41586-020-2312-y.

15. Dalkara, T.; Gursoy-Ozdemir, Y.; Yemisci, M. Brain microvascular pericytes in health and disease. Acta neuropathologica 2011, 122, 1–9, doi:10.1007/s00401-011-0847-6.

16. Consortium, G.T.; Laboratory, D.A.; Coordinating Center -Analysis Working G.; Statistical Methods groups-Analysis Working, G.; Enhancing, G.g.; Fund, N.I.H.C.; Nih/Nci; Nih/Nhgri; Nih/Nimh; Nih/Nida, et al. Genetic effects on gene expression across human tissues. Nature 2017, 550, 204–213, doi:10.1038/nature24277.

17. Bertrand, L.; Cho, H.J.; Toborek, M. Blood–brain barrier pericytes as a target for HIV-1 infection. Brain 2019, 142, 502–511, doi:10.1093/brain/awy339.

18. Brann, D.H.; Tsukahara, T.; Weinreb, C.; Lipovsek, M.; Van den Berge, K.; Gong, B.; Chance, R.; Macaulay, I.C.; Chou, H.-j.; Fletcher, R., et al. Non-neuronal expression of SARS-CoV-2 entry genes in the olfactory system suggests mechanisms underlying COVID-19-associated anosmia. bioRxiv 2020, 10.1101/2020.03.25.009084, 2020.2003.2025.009084, doi:10.1101/2020.03.25.009084.

19. Fodoulian, L.; Tuberosa, J.; Rossier, D.; Landis, B.N.; Carleton, A.; Rodriguez, I. SARS-CoV-2 receptor and entry genes are expressed by sustentacular cells in the human olfactory neuroepithelium. bioRxiv 2020, 10.1101/2020.03.31.013268, 2020.2003.2031.013268, doi:10.1101/2020.03.31.013268.

20. Muus, C.; Luecken, M.D.; Eraslan, G.; Waghray, A.; Heimberg, G.; Sikkema, L.; Kobayashi, Y.; Vaishnav, E.D.; Subramanian, A.; Smilie, C., et al. Integrated analyses of single-cell atlases reveal age, gender, and smoking status associations with cell type-specific expression of mediators of SARS-CoV-2 viral entry and highlights inflammatory programs in putative target cells. bioRxiv 2020, 10.1101/2020.04.19.049254, 2020.2004.2019.049254, doi:10.1101/2020.04.19.049254.

21. Chen, R.; Wang, K.; Yu, J.; Chen, Z.; Wen, C.; Xu, Z. The spatial and cell-type distribution of SARS-CoV-2 receptor ACE2 in human and mouse brain. bioRxiv 2020, 10.1101/2020.04.07.030650, 2020.2004.2007.030650, doi:10.1101/2020.04.07.030650.

22. Yang, L.; Han, Y.; Nilsson-Payant, B.E.; Gupta, V.; Wang, P.; Duan, X.; Tang, X.; Zhu, J.; Zhao, Z.; Jaffre, F., et al. A Human Pluripotent Stem Cell-based Platform to Study SARS-CoV-2 Tropism and Model Virus Infection in Human Cells and Organoids. Cell stem cell 2020, 27, 125–136 e127, doi:10.1016/j.stem.2020.06.015.

23. Bullen, C.K.; Hogberg, H.T.; Bahadirli-Talbott, A.; Bishai, W.R.; Hartung, T.; Keuthan, C.; Looney, M.M.; Pekosz, A.; Romero, J.C.; Sille, F.C.M., et al. Infectability of human BrainSphere neurons suggests neurotropism of SARS-CoV-2. Altex 2020, 10.14573/altex.2006111, doi:10.14573/altex.2006111.

24. Storey, J.D.; Bass, A.J.; Dabney, A.; Robinson, D. qvalue: Q-value estimation for false discovery rate control. R package version 2.20.0 2020, http://github.com/jdstorey/qvalue.

25. Durinck, S.; Spellman, P.T.; Birney, E.; Huber, W. Mapping identifiers for the integration of genomic datasets with the R/Bioconductor package biomaRt. Nature protocols 2009, 4, 1184–1191, doi:10.1038/nprot.2009.97.

26. Subramanian, A.; Tamayo, P.; Mootha, V.K.; Mukherjee, S.; Ebert, B.L.; Gillette, M.A.; Paulovich, A.; Pomeroy, S.L.; Golub, T.R.; Lander, E.S., et al. Gene set enrichment analysis: a knowledge-based approach for interpreting genome-wide expression profiles. Proceedings of the National Academy of Sciences of the United States of America 2005, 102, 15545–15550, doi:10.1073/pnas.0506580102.

